# Benchmarking AI Models for *In Silico* Gene Perturbation of Cells

**DOI:** 10.1101/2024.12.20.629581

**Authors:** Chen Li, Haoxiang Gao, Yuli She, Haiyang Bian, Qing Chen, Kai Liu, Lei Wei, Xuegong Zhang

**Affiliations:** MOE Key Laboratory of Bioinformatics and Bioinformatics Division of BNRIST, Department of Automation, Tsinghua University, Beijing 100084, China; Anew Therapeutics Pte. Ltd., Singapore 068896, Singapore; Center for Synthetic and Systems Biology, School of Life Sciences and School of Medicine, Tsinghua University, Beijing 100084, China

## Abstract

Understanding perturbations at the single-cell level is essential for unraveling cellular mechanisms and their implications in health and disease. The growing availability of biological data has driven the development of a variety of *in silico* perturbation methods designed for single-cell analysis, which offer a means to address many inherent limitations of experimental approaches. However, these computational methods are often tailored to specific scenarios and validated on limited datasets and metrics, making their evaluation and comparison challenging. In this work, we introduce a comprehensive benchmarking framework to systematically evaluate *in silico* perturbation methods across four key scenarios: predicting effects of unseen perturbations in known cell types, predicting effects of observed perturbations in unseen cell types, zero-shot transfer to bulk RNA-seq of cell lines, and application to real-world biological cases. For each scenario, we curated diverse and abundant datasets, standardizing them into flexible formats to enable efficient analysis. Additionally, we developed multiple metrics tailored to each scenario, facilitating a thorough and comparative evaluation of these methods. Our benchmarking study assessed 10 methods, ranging from linear baselines to advanced machine learning approaches, across these scenarios. While some methods demonstrated surprising efficacy in specific contexts, significant challenges remain, particularly in zero-shot predictions and the modeling of complex biological processes. This work provides a valuable resource for evaluating and improving *in silico* perturbation methods, serving as a foundation for bridging computational predictions with experimental validation and real-world biological applications.

## Introduction

Understanding responses of gene expression to gene perturbations in cells is crucial for decoding cellular functions and has important implications for many applications in biomedical research. With advancements in single-cell sequencing and gene perturbation technologies, methods such as Perturb-seq^1^ and CROP-seq^2^, which integrate single-cell RNA sequencing (scRNA-seq) with CRISPR-based perturbations, enable large-scale pooled assays^3, 4^ to investigate gene functions and cellular responses at single-cell resolution. However, experimental approaches face significant limitations: with ∼20,000 protein-coding genes and ∼400 major cell types in humans^5^, it is practically impossible to explore all gene perturbations across all cell types, let alone explore the vast array of gene combinations. Moreover, these experiments usually require multi-day cell incubation, but many cell types cannot survive in culture for extended periods, limiting its applicability to many biological contexts.

These constraints have spurred the development of a number of artificial intelligence (AI) models for *in silico* gene perturbation, which predict cellular states (typically represented as gene expression profiles) following gene perturbations. Dynamo^6^ introduced the concept of *in silico* perturbation for single-cell datasets. It leverages the analytical Jacobian of a reconstructed vector field function to encode gene regulatory networks (GRNs). By propagating perturbations through the GRN, Dynamo predicts cell-fate outcomes following perturbation. CellOracle^7^ developed an algorithm to infer causal GRNs using both scRNA-seq and single-cell assay for transposase-accessible chromatin sequencing (scATAC-seq) data, enabling GRN-based *in silico* perturbations. GEARS^8^ marked a significant advancement as the first method capable of predicting transcriptional outcomes for unseen perturbations by training on Perturb-seq data, learning gene embeddings through a gene co-expression graph and perturbation embeddings through a Gene Ontology (GO) term knowledge graph. Inspired by GEARS, subsequent methods^9, 10^ refined this strategy to achieve improved performance. In parallel, approaches focusing on predicting specific treatment responses in novel cell types have emerged, with the variational auto-encoder (VAE) framework widely adopted. Examples include scGen^11^, CPA^12^, sVAE^13^, and SAMS-VAE^14^. Additionally, optimal transport theory has also been applied to *in silico* perturbation, with notable examples including CellOT^15^ and CINEMA-OT^16^. Recently, single-cell foundation models have introduced a new paradigm in *in silico* perturbation. Models like scGPT^17^ and scFoundation^18^ leverage large-scale pretrained architectures to deliver promising performance. Overall, these *in silico* gene perturbation methods utilized advanced statistical and deep learning algorithms, showing great potential to expand our understanding of gene perturbations and cellular responses for various cell types.

However, several challenges hinder the development and comparison of *in silico* gene perturbation methods. These methods are often tailored to specific scenarios, validated on limited datasets, and assessed using inconsistent metrics. For example, GEARS aims to predict unseen perturbations in known cell types, whereas scGen and CPA focus on predicting responses in unseen cell types with known perturbations. In addition, these methods are typically evaluated on narrowly focused datasets and metrics suited to specific contexts. For instance, Dynamo and CellOracle assessed their performance on datasets representing biological developmental trajectories, such as the hematopoietic data, while GEARS and scGPT used single-cell perturbation datasets for validation. This reliance on limited and narrowly focused datasets introduces biases into the evaluation process, complicating the assessment of their effectiveness and generalizability. Furthermore, the inconsistency in evaluation metrics across studies, ranging from qualitative metrics like trajectory direction alignment to various quantitative metrics, further complicates performance comparisons.

Some benchmarking studies have attempted to evaluate existing *in silico* perturbation methods^19–22^, but they often focus on a limited subset of methods, datasets, or scenarios, leading to incomplete or biased conclusions. For example, Wu et al.^20^ proposed a perturbation benchmark for various machine learning methods, but it did not include evaluations for predicting unseen perturbations. Similarly, Eltze el al.^22^ conducted a study claiming that existing deep learning methods do not outperform simple linear models, but the evaluation relied on the L2 error across all genes, which may focus more on the significantly varied genes while overlooking the invariant ones. These limitations highlight the absence of a comprehensive benchmarking framework that provides standardized datasets and evaluation metrics to fairly compare methods. Just as the MNIST^23^ and ImageNet^24^ datasets revolutionized computer vision by enabling systematic performance comparisons, a well-designed benchmark could play a transformative role in the development of *in silico* perturbation methods.

In this work, we presented a comprehensive benchmarking framework with rich collection of data to systematically evaluate *in silico* gene perturbation method. Depending on various usage requirements, we categorized *in silico* perturbation methods into four distinct scenarios: unseen perturbation transfer, useen cell type transfer, zero-shot transfer, and cell state transition prediction. We curated and filtered the gene perturbation datasets of scPerturb, retaining 17 single-cell datasets encompassing 984,000 cells and 3,190 perturbations. We designed a comprehensive set of evaluation metrics to assess various aspects of method performance. We conducted benchmarking study to assess 10 methods across all scenarios, including linear baseline methods, specific machine learning methods and large pretrained AI models. We observed that while some methods demonstrated surprising efficacy in specific contexts, significant challenges remain, particularly in zero-shot predictions and the modeling of complex biological processes. The presented benchmarking framework and datasets offered a valuable resource for the scientific community, serving as a foundation to promote the further development and refinement of AI models for *in silico* perturbation.

## Results

### Framework for benchmarking *in silico* perturbation

We present a benchmarking framework designed to evaluate *in silico* gene perturbation methods across four biologically and computationally relevant scenarios (**Fig. 1a**). The unseen perturbation transfer scenario assesses the ability to predict effects of previously unobserved perturbations in known cell types. The unseen cell type transfer scenario evaluates the prediction of known perturbations in previously unobserved cell types. The zero-shot transfer scenario evaluates model performance in predicting perturbation effects measured within the CMAP framework, where perturbations are assessed using bulk RNA-seq data of cell lines. The cell state transition prediction scenario focuses on predicting effects of key genes driving cell state transitions in specific biological processes. The first two scenarios emphasize well-defined transfer tasks with available training data, while the latter two reflect more challenging real-world applications where *in silico* perturbation tools often need to generalize without task-specific training.

**Fig. 1.**
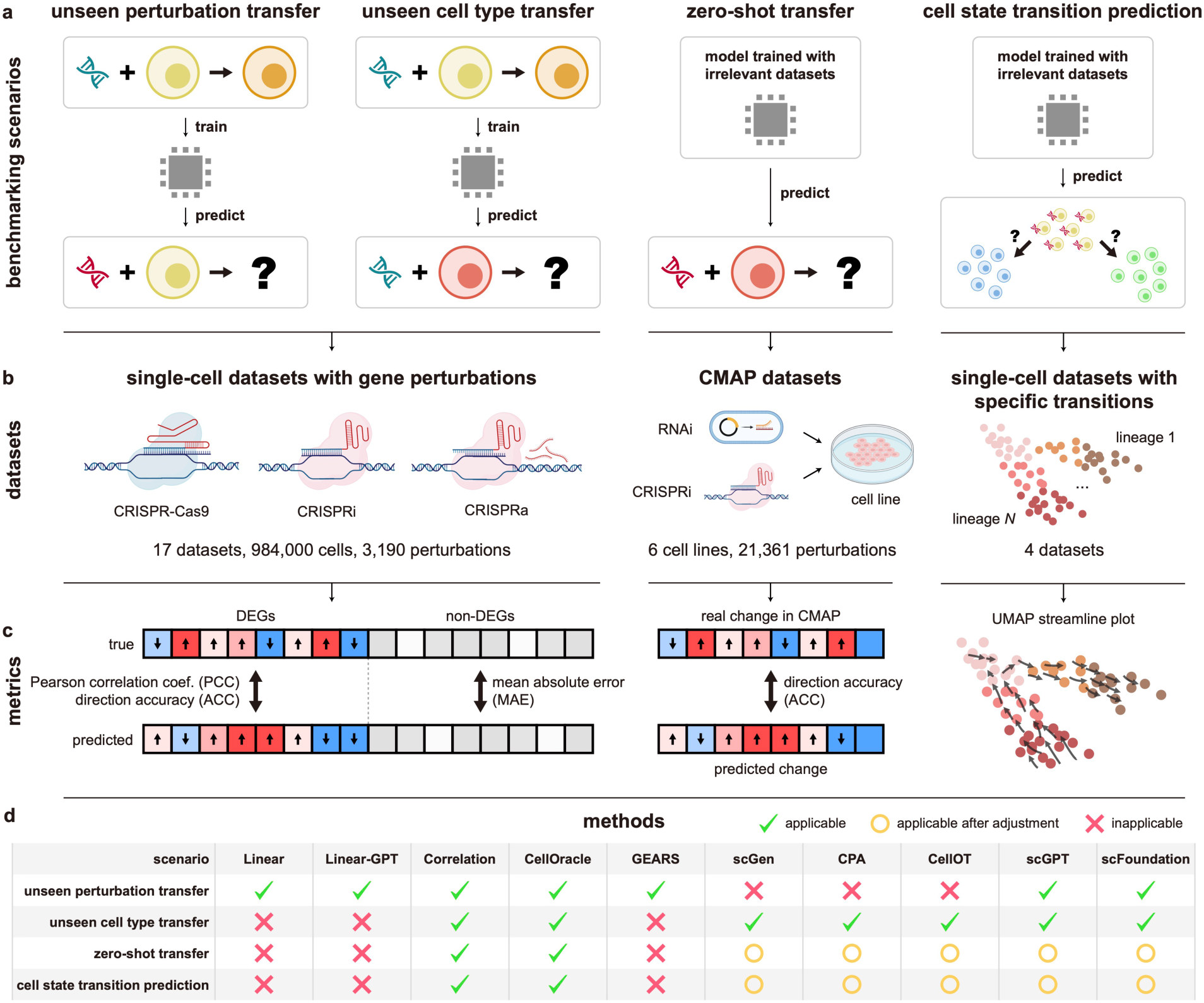
Overview of the *in silico* perturbation benchmarking framework. **(a)** Four benchmarking scenarios evaluated in this study. **(b)** Datasets curated for benchmarking. **(c)** Metrics for designed for evaluation across scenarios. **(d)** Methods benchmarked in this study.

For the unseen perturbation transfer scenario, we collected single-cell perturbation datasets from scPerturb^25^ as benchmarking data (**Fig. 1b**). After quality control, preprocessing, and reformatting, we curated 17 single-cell datasets, resulting in 984,000 cells and 3,190 gene perturbations, including CRISPR-Cas9 knockout, CRISPR interference (CRISPRi), and CRISPR activation (CRISPRa) (Methods). Genes were classified into differentially expressed genes (DEGs) and non-DEGs. For DEGs, benchmarking metrics included Pearson correlation coefficient (PCC) and direction accuracy (ACC), while performance for non-DEGs was assessed using mean absolute error (MAE) (**Fig. 1c**).

The unseen cell type transfer scenario used two datasets from Replogle et al.^4^, which contained the largest number of perturbations among our curated datasets, selecting 490 shared perturbations as benchmarking data (**Fig. 1b**). The dataset from the K562 cell line was used for training, while the RPE1 dataset served as testing data. Evaluation metrics were consistent with those in the unseen perturbation transfer scenario (**Fig. 1c**).

In the zero-shot transfer scenario, we employed Connectivity Map (CMAP) data^26^, measured at the bulk level, to evaluate model performance (**Fig. 1b**). Single-cell RNA-seq datasets^27, 28^ from six cell lines overlapping with CMAP data were used as the input for *in silico* perturbation methods. After quality control, a total of 21,361 perturbations from six cell lines served as the ground truth (Methods). For benchmarking, we selected the top significant genes (genes with high absolute z-scores) from the CMAP data for each perturbation and calculated direction accuracy (ACC) of these genes (**Fig. 1c**, Methods).

For the cell state transition prediction scenario, we collected four scRNA-seq datasets^29–32^ representing distinct biological processes, including development, reprogramming, and disease progression (**Fig. 1b**). For each dataset, there are substantial reliable evidence showing that individual genes or specific gene combinations can drive certain cell state transitions. Qualitative evaluation included streamline plots in uniform manifold approximation and projection (UMAP) space to visualize predictions, while a ranking transition score (RTS) was introduced to quantitatively measure accuracy of predicted transitions in response to perturbations (**Fig. 1c**, Methods).

Considering reproducibility and broad applicability, we evaluated 10 methods in this work, spanning linear models, gene regulatory network (GRN)-based approaches, and machine learning techniques (**Fig. 1d**, Methods). These included linear models^22^ (with and without scGPT embeddings, referred to as Linear and Linear-GPT, respectively), correlation-based method, CellOracle^7^, GEARS^8^, scGen^11^, CPA^12^, CellOT^15^, scGPT^17^, and scFoundation^18^. Given the varying design and applicability of these methods, we selected and evaluated an appropriate subset for each specific scenario.

### Benchmarking in the unseen perturbation transfer scenario

In the unseen perturbation transfer scenario (**Fig. 2a**), we utilized version 1.3 of the scPerturb database as our benchmarking data. scPerturb contains 50 single-cell perturbation datasets for RNA and protein expression. Following rigorous data filtering and standardized processing (Methods), we curated 17 scRNA-seq datasets^1–4, 33–37^ for analysis.

**Fig. 2.**
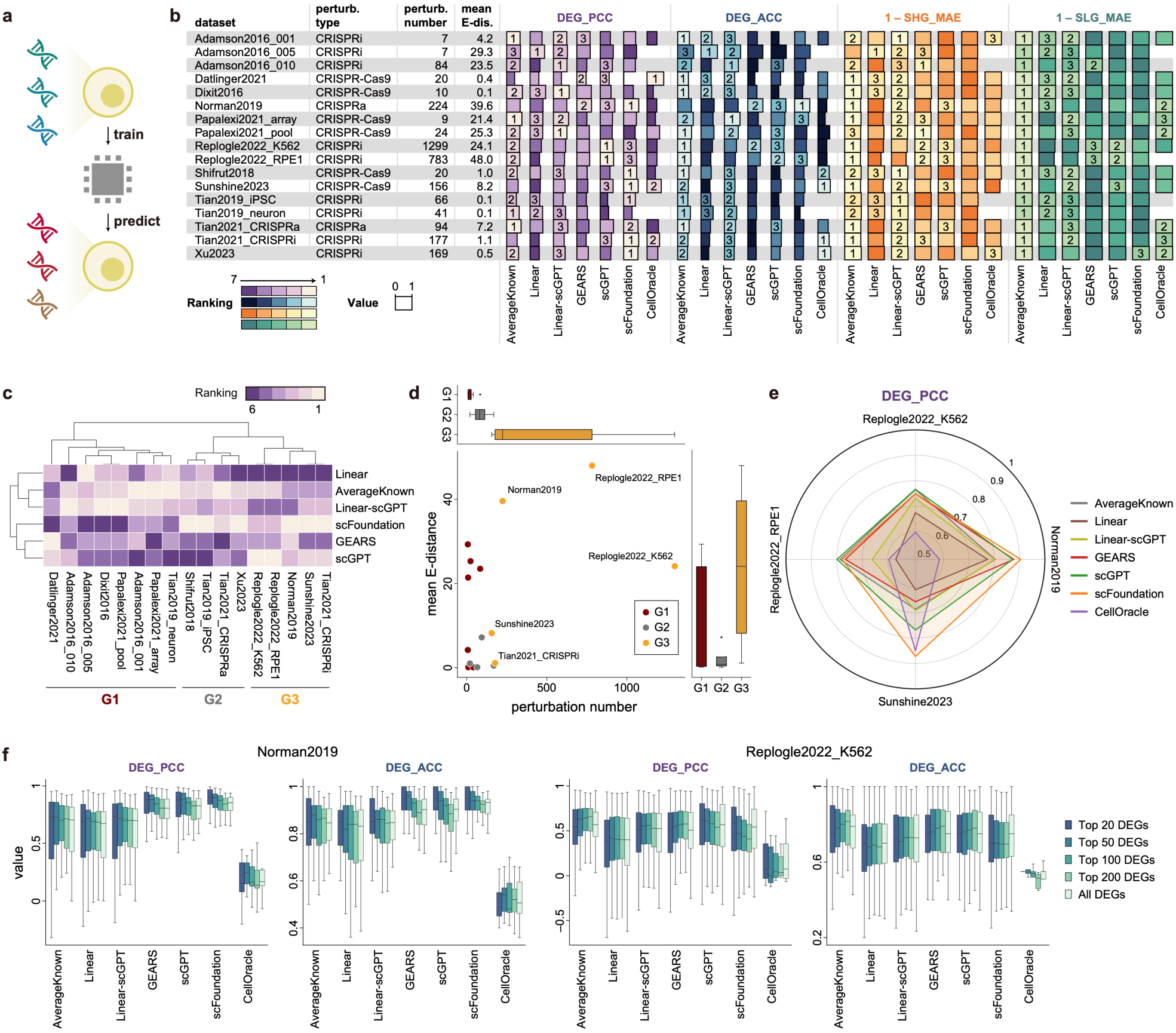
Benchmarking in the unseen perturbation transfer scenario. **(a),** Schematic diagram of unseen perturbation transfer. **(b)** Performance of methods on 17 single-cell perturbation datasets. Each dataset is exhibited with attributes of perturbation type, perturbation number, and mean E-distance. Metrics are divided into 4 parts: DEG_PCC (Pearson correlation coefficient for differentially expressed genes), DEG_ACC (direction accuracy for differentially expressed genes), SHG_MAE (mean absolute error for the genes that are stably highly expressed), and SLG_MAE (mean absolute error for the genes that are stably lowly expressed). The top three methods for each metric on each dataset are highlighted. Color indicates ranking, and bar length represents the magnitude of the metric. For DEG_PCC, negative values are not shown. **(c)** Hierarchically-clustered heatmap of methods’ performance rankings across datasets. The datasets are broadly clustered into 3 groups (G1, G2, and G3). **(d)** Scatter plot showing the mean E-distance and the number of perturbations for each dataset, accompanied by bar plots comparing the mean E-distance and perturbation number across different dataset groups. **(e)** Radar plot of DEG_PCC for 7 methods across 4 selected datasets. **(f)** Box plots comparing DEG_PCC and DEG_ACC of 7 methods on Norman2019 and Replogle2022_K562 datasets, with different levels of DEGs (top 20, 50, 100, 200, and all) selected for each metric. Center line, median; box limits, upper and lower quartiles; whiskers, 1.5× interquartile range.

We calculated the E-distance, a statistical measure of the distance between point clouds, applied to pre-and post-perturbation cell pairs for each perturbation as an indicator of the magnitude of perturbation effect^25^. The number of perturbations in these 17 datasets varies widely, ranging from 7 (Adamson2016_001 and Adamson2016_005) to 1,299 (Replogle2022_K562), highlighting substantial differences in data scale. Similarly, the mean E-distances also vary considerably across datasets, spanning from 0.1 to 48.0, likely reflecting differences in experimental conditions. We summarized key attributes of these datasets, including the gene perturbation technique, the number of perturbations and the mean E-distance in **Fig. 2b**.

Considering that only a subset of genes is affected by specific perturbations, we categorized the genes in the gene expression profiles for each perturbation into three sets: Differentially expressed genes (DEG), which exhibit significant expression differences between pre-and post-perturbation cells; (2) Stably highly expressed genes (SHG), which are not differentially expressed and show high expression in pre-perturbation cells; and (3) Stably lowly expressed genes (SLG), which are not differentially expressed and show low expression in pre-perturbation cells. For the DEG set, we evaluated performance using PCC (DEG_PCC) and ACC (DEG_ACC) values, while for the SHG and SLG set, we assessed performance using MAE values, referred to as SHG_MAE and SLG_MAE, respectively (Methods).

To benchmark the methods, we evaluated the performance of six methods: Linear, Linear-GPT, CellOracle, GEARS, scGPT, and scFoundation (Methods). We also included a basic approach that averages gene expression across all cells within all known perturbations as the prediction of unseen perturbations (referred to as KnownAverage).

The benchmarking results across the 17 datasets were summarized in **Fig. 2b**. Notably, the KnownAverage method consistently demonstrated some of the best overall performance across all four types of metrics. Specifically, for the DEG_PCC metric, the KnownAverage method ranked among the top three in most datasets. The Linear and Linear-scGPT method also achieved top ranks in specific datasets, such as Adamson2016_005, Dixit2016, and Papalexi2021_array. Meanwhile, single-cell foundation models, including scGPT and scFoundation, outperformed other methods in datasets such as Replogle2022_K562, Replogle2022_RPE1, and Norman2019. GEARS and CellOracle achieved top ranking in only a few datasets, with CellOracle showing the weakest overall performance. For DEG_ACC, the overall performance trends mirrored those of DEG_PCC, with the KnownAverage method exhibiting superior performance and ranking top in nearly all the datasets.

For both SHG_MAE and SLG_MAE, the KnownAverage method again achieved best performance in nearly all datasets, and Linear-scGPT consistently ranks in the top three for most cases. Although deep learning methods such as GEARS, scGPT, and scFoundation did not consistently achieve top ranks, their metrics were not significantly worse than those of the best-performing methods. The Linear method exhibited a notable performance drop in specific datasets, such as Papalexi2021_array and Replogle2022_RPE1.

We performed hierarchical clustering on the heatmap of DEG_PCC rankings of models across the 17 datasets (we did not include CellOracle because it is not compatible with all datasets.). The results revealed distinct performance patterns, grouping the datasets into three clusters: G1, G2, and G3 (**Fig. 2c**). To better understand these groupings, we examined the perturbation number and mean E-distance for each dataset and compared the average attributes of the three groups (**Fig. 2d**). Group G3 exhibited the highest average perturbation number (527.8) and mean E-distance (24.2), indicating that these datasets include a wide range of perturbations with significant phenotypic effects pre-perturbation cells. Single-cell foundation models, such as scGPT and scFoundation, performed best on G3 datasets, likely due to the abundance of heterogeneous perturbations in the training data, which supports the learning capabilities of large deep learning models.

Group G2 displayed a moderate average perturbation number (87.2) but a low average E-distance (2.2), suggesting that perturbations in these datasets induced only subtle shifts in expression profiles. Interestingly, scFoundation achieved the best performance on G2 datasets, demonstrating its ability to capture slight perturbation effects more effectively than other methods, including scGPT.

Group G1 showed the lowest average perturbation number (25.2). In these datasets, simple methods like KnownAverage, Linear, and Linear-scGPT outperformed deep learning methods. This performance gap is likely attributed to the restricted number of perturbations, which limits the capacity of deep learning models to effectively learn perturbation effects. Additionally, some datasets in G1 include perturbed genes that belong to the same functional group. For instance, in the Datlinger2021 dataset^2^, the 20 perturbed genes were specifically selected for their relevance to the T cell receptor pathway. This likely reduces the complexity and heterogeneity of the perturbations, making the simpler methods more effective in modeling the observed perturbation effects.

Based on these observations, we conclude that datasets with higher perturbation numbers and larger E-distances, such as those in D3, are better suited for extending to genome-wide unseen perturbations. These characteristics provide a robust foundation for fair and comprehensive comparisons of different methods.

We further selected 4 datasets from Group G3 for final benchmarking, focusing on datasets with a high number of perturbations (>150) and substantial E-distances (>5): Replogle2022_K562^4^, Replogle2022_RPE1^4^, Norman2019^38^, and Sunshine2023^39^. For the DEG_PCC metric, single-cell foundation models, including scFoundation and scGPT, outperformed all other methods, indicating their ability to capture significant gene expression changes (**Fig. 2e**). Overall, scFoundation outperformed scGPT, particularly in the Sunshine2023 dataset. The AverageKnown method and GEARS exhibited comparable performance, with average PCCs of 0.734 and 0.722, respectively. The Linear method and CellOracle showed generally poor performance across these datasets. Though Linear-scGPT outperformed the Linear method, it still did not surpass the AverageKnown method, raising questions about whether the gene embeddings from scGPT genuinely aids this task. For the DEG_ACC metric, the AverageKnown method, scGPT, and scFoundation demonstrated comparable performance, with average ACCs of 0.732, 0.726, and 0.705, respectively (**Fig. S1**). Other methods, including GEARS, Linear-scGPT, Linear, and CellOracle, showed lower ACCs of 0.692, 0.689, 0.621, and 0.601, respectively, indicating limited effectiveness in predicting the direction of gene expression changes. Additionally, single-cell foundation models demonstrated improved performance when focusing on higher-ranked DEGs, as shown in **Fig. 2f** and **S2**, suggesting that these models are better at capturing the most significant transcriptional changes.

For the SHG_MAE and SLG_MAE metrics, the mean method, scGPT, and scFoundation consistently performed well (**Fig. S1**). CellOracle performed slightly worse on the Sunshine2023 dataset for SHG_MAE, while GEARS showed slightly lower performance on both the Sunshine2023 and Norman2019 datasets for SLG_MAE. In contrast, linear-type methods performed poorly across both SHG_MAE and SLG_MAE.

Overall, single-cell foundation models, such as scGPT and scFoundation, emerged as the best-performing methods in the unseen perturbation transfer scenario. With sufficient perturbation data for training, they are capable of accurately predicting the expression changes of DEGs while preserving the expression levels of non-DEGs. The results suggest that single-cell foundation models could be effective choices for extending predictions to unseen perturbations. Interestingly, the simplest approach, AverageKnown, also performed remarkably well across various datasets, including large ones like Replogle2022_K562. By visualizing the average expression changes of perturbations in this dataset using a clustering heatmap, we observed that more than half of the perturbed genes were grouped into a single cluster (**Fig. S3**). As a result, the AverageKnown method can achieve good performance by simply averaging these perturbed cells. This raises the concern that simple methods may unexpectedly perform well due to biases, and researchers should exercise caution when applying them. GEARS also demonstrated potential as an advanced solution for unseen perturbations but performed slightly worse than the single-cell foundation models. In contrast, Linear, Linear-scGPT, and CellOracle consistently underperformed compared to deep learning-based methods.

### Benchmarking in the unseen cell type transfer scenario

In the unseen cell type transfer scenario (**Fig. 3a**), we utilized the Replogle2022_K562 and Replogle2022_RPE1 datasets from the same study^4^ for benchmarking. These datasets were generated using Perturb-seq experiments conducted on two distinct cell lines: K562, a lymphoblast cell line derived from a patient with chronic myelogenous leukemia, and RPE1, a retinal pigment epithelial cell line. After standardized data processing, we identified 490 shared gene perturbations and 3,144 overlapping genes between the two datasets.

**Fig. 3.**
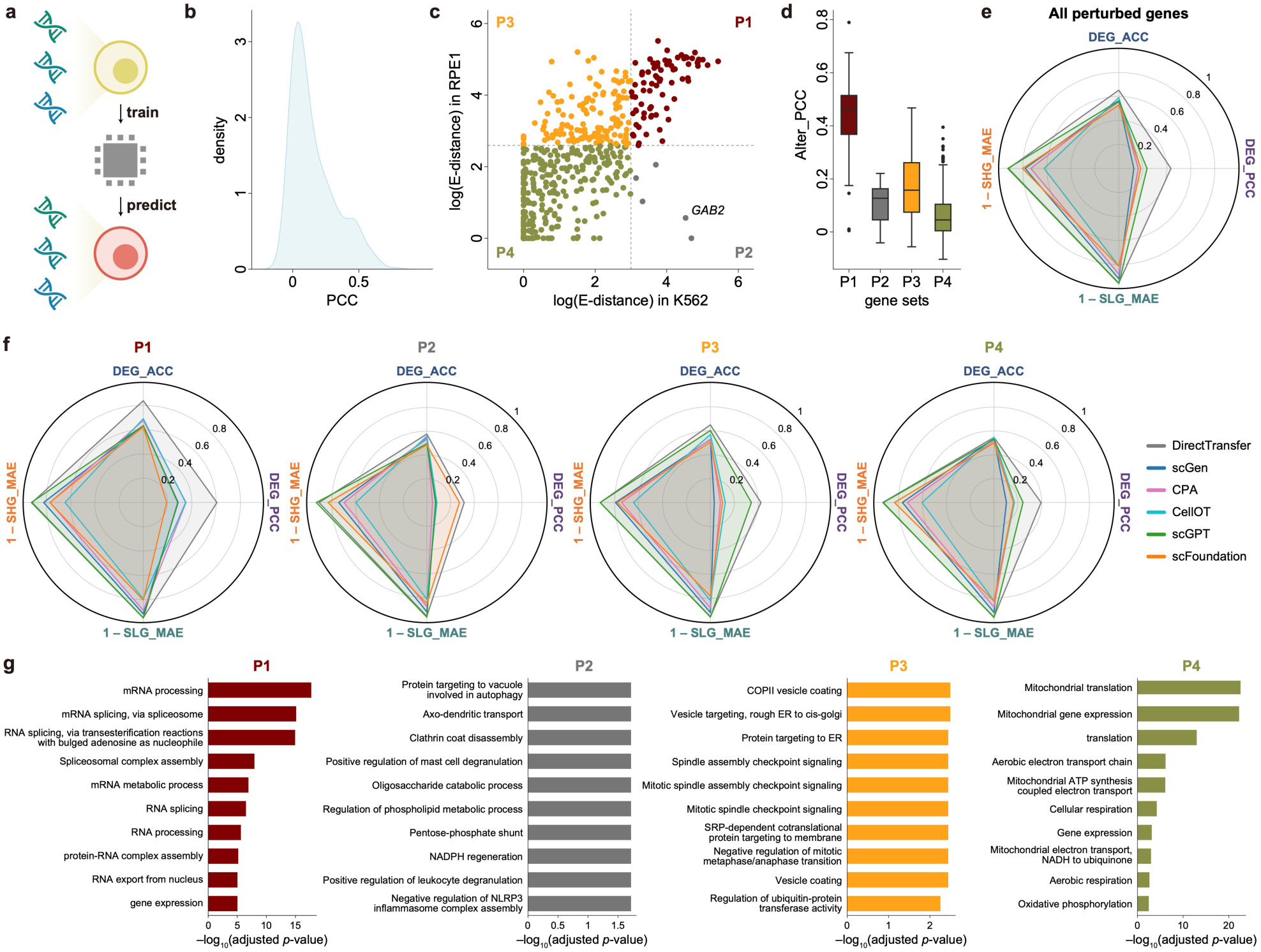
Benchmarking in the unseen cell type transfer scenario. **(a)** Schematic diagram of unseen cell type transfer. **(b)** Density plot illustrating the distribution of PCCs of gene expression alterations across 490 common perturbations. **(c)** Scatter plots of 490 common perturbations, with the x-axis and y-axis represent log(E-distance) in the K562 and RPE1 cell lines, respectively. The perturbations are grouped into four categories (P1, P2, P3, and P4) based on log(E-distance) thresholds. **(d)** Box plots showing the PCCs of gene expression alterations (Alter_PCC) across different perturbation groups. Center line, median; box limits, upper and lower quartiles; whiskers, 1.5× interquartile range. **(e)** Radar plot comparing DEG_PCC, DEG_ACC, SHG_MAE, and SLG_MAE across the 490 perturbations for 6 methods. **(f)** Radar plots comparing DEG_PCC for 6 methods across 4 perturbation groups. **(g)** GO enrichment analysis for genes in each perturbation group.

For benchmarking, all models were trained on the shared perturbations from Replogle2022_K562 and then tested on the corresponding perturbations in Replogle2022_RPE1. Metrics were calculated based on the 3,144 overlapping genes (Methods). In this scenario, we selected scGen, CPA, CellOT, scGPT, and scFoundation as the benchmarking methods. Additionally, we included a simple baseline method, DirectTransfer, which directly transfers the gene expression changes of perturbations from Replogle2022_K562 to Replogle2022_RPE1 without additional model adjustments.

Before comparing different methods, we first assessed the similarity of shared perturbations between the two datasets. For each perturbation in each dataset, we calculated the gene expression alterations of averaged gene expression profiles between perturbed cells and unperturbed cells, and then calculated the PCCs of gene expression alterations (Alter_PCCs) for each perturbation in Replogle2022_K562 and Replogle2022_RPE1. The similarity provides a critical indicator of the difficulty in unseen cell type transfer prediction. As shown in **Fig. 3b**, the distribution of Alter_PCCs exhibits a bimodal pattern: a large group of perturbations clusters around 0.05, indicating divergent effects on gene expression between the two cell lines, while a smaller group clusters around 0.45, suggesting potentially similar effects. The distributions of Alter_PCC values suggest that these perturbations should be categorized into distinct groups for a more comprehensive benchmarking.

We further visualized each perturbation’s E-distances in Replogle2022_K562 and Replogle2022_RPE1 (**Fig. 3c**). Using a log(E-distance) threshold of 3 in each dataset, we classified the 490 shared perturbations into four sets (Fig. 3C): P1 (above the threshold in both datasets), P2 (above in K562 but below in RPE1), P3 (below in K562 but above in RPE1), and P4 (below in both datasets). These classifications reflect the varying effects of perturbations, which are associated with differing Alter_PCC values. For instance, P1 comprises perturbations that induce substantial and similar phenotypic changes in both K562 and RPE1 cell lines, with a median Alter_PCC value of 0.46—significantly higher than those observed in the other three sets (**Fig. 3d**).

We first evaluated all methods across overall 490 shared perturbations (**Fig. 3e**). Results showed that DirectTransfer outperforms all other methods across various metrics, including DEG_PCC, DEG_ACC, SHG_MAE, and SLG_MAE. For example, the mean DEG_PCC of DirectTransfer was 0.606, significantly higher than the other five methods.

Notably, DirectTransfer achieved its highest performance in the P1 group (**Fig. 3f**). CellOT and CPA demonstrated moderate performance in DEG_PCC and DEG_ACC, and CellOT showed a decline in SHG_MAE and SLG_MAE. In contrast, scGPT, scFoundation, and scGen demonstrated poor performance overall. Gene ontology (GO) enrichment analysis revealed that P1 perturbations are associated with fundamental biological processes such as mRNA processing and RNA splicing (Fig. 3G), explaining their significant and consistent effects across K562 and RPE1 cell lines. This indicates that predicting the effects of P1 perturbations in unseen cell types is relatively easier for current methods.

Perturbations in P2 were associated with only five genes, such as *GAB2*, which is involved in the positive regulation of leukocyte and mast cell degranulation (**Fig. 3g**), and this suggests genes in P2 may play an important role in K562, but have a weak effect in RPE1. For perturbations in P2, scFoundation achieved performance close to DirectTransfer in DEG_PCC, while scGen, CPA, CellOT, and scGPT yielded extremely low DEG_PCC values (**Fig. 3f**), indicating that these models struggle to quantitatively capture the expression changes of DEGs.

Perturbations in P3 were enriched in some biological processes related to protein production and secretion, such as COPII vesicle coating and protein targeting to ER (**Fig. 3g**). This likely reflects the epithelial nature of RPE1 cells, which are involved in more processing and transport of membrane and secretory proteins. On P3 perturbations, scGPT performed comparably to DirectTransfer across all metrics, whereas other methods showed notably poorer performance (**Fig. 3f**).

Perturbations in P4 corresponded to genes with slight effects across both K562 and RPE1 cells. These perturbations were enriched in some biological processes that are not predominant in cells, such as mitochondrial translation and mitochondrial gene expression (**Fig. 3g**). DirectTransfer still maintained the best overall performance on P4 perturbations, followed by scGPT and scFoundation (**Fig. 3f**).

All the above results showed that advanced machine learning or statistical methods failed to outperform the simplest baseline, DirectTransfer, in the unseen cell type transfer scenario. While some methods have demonstrated effectiveness in predicting specific treatment responses in unseen cell types, such as scGen predicting the cellular response to IFN-β stimulation in CD4 T cells^11^, and CellOT predicting responses of patients with glioblastoma to panobinostat^15^, these models struggled in the context of gene perturbations. Transformer-based models, such as scGPT and scFoundation, generally outperformed autoencoder-based methods, such as scGen, CPA, and CellOT, but still fell significantly short of the baseline performance of DirectTransfer. But DirectTransfer completely ignores the specificity of different cell types, making it unsuitable for practical use. This highlights the ongoing need for new, more effective methods tailored to the unseen cell type transfer scenario. Furthermore, when applying *in silico* perturbation methods to predict effects on unseen cell types, the choice of gene perturbations plays a crucial role in determining the results. Perturbing certain housekeeping genes may elicit similar cellular responses across different cell types, as seen in the P1 and P4 groups, while other genes may have varying functional importance across cell types, as in P2 and P3, depending on their specific roles across different cell types.

### Benchmarking in the zero-shot transfer scenario

In practical applications, researchers often aim to apply *in silico* perturbation methods directly to their own datasets, where training data from matching experimental settings with perturbations are typically unavailable. To address this scenario, we established a benchmark for evaluating model performance in transferring to bulk RNA-seq data (**Fig. 4a**).

**Fig. 4.**
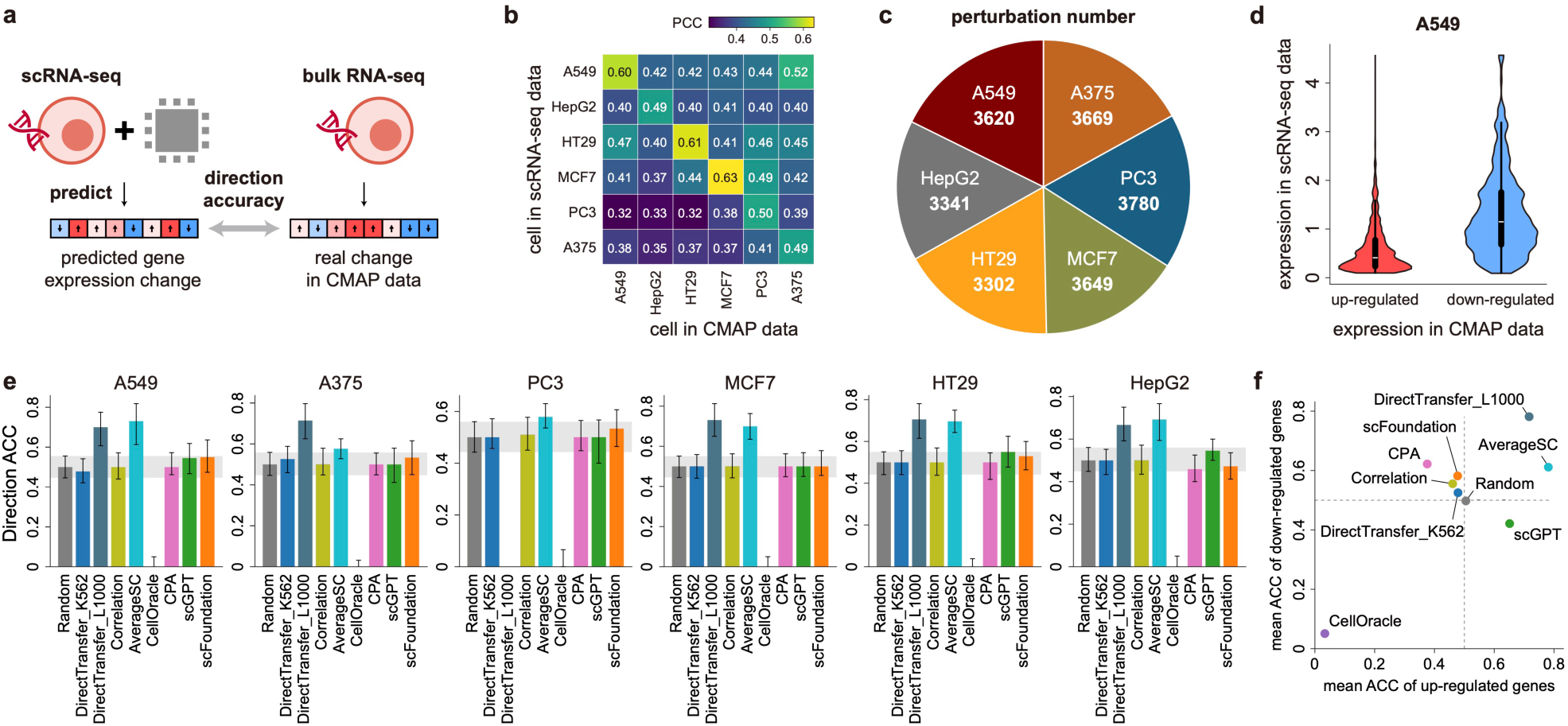
Benchmarking in the zero-shot transfer scenario. **(a)** Schematic diagram of zero-shot transfer. **(b)** Heatmap of correlations between single-cell data and CMAP data. **(c)** Pie chart illustrating the distribution of perturbations across six cell lines in the CMAP dataset. **(d)** Expression levels of CMAP’s up-regulated and down-regulated genes in single-cell data. Up-regulated genes: genes that are up-regulated after perturbations in CMAP data; down-regulated genes: genes that are down-regulated after perturbations in CMAP data. **(e)** Bar plots comparing direction accuracy (ACC) across 9 methods for the 6 cell lines. Each bar represents the median ACC, with black bars indicating the upper and lower quartiles. The range of gray bars represent the upper quartile and lower quartile of the performance of Random. **(f)** Scatter plots showing the mean ACC of up-regulated and down-regulated genes. The gray dashed lines indicate the performance of Random for up-regulated and down-regulated genes.

We used CMAP data as the testing data for benchmarking. We collected single-cell datasets^27, 28^ of six cell lines that overlap with CMAP data: A375, PC3, MCF7, HT29, HepG2, and A549. Consistency between gene expression profiles in the single-cell data and the corresponding CMAP data was verified (**Fig. 4b**, Methods). These single-cell datasets were processed as inputs to *in silico* methods to directly predict gene expression changes resulting from gene perturbations without any additional training. For each perturbation, we selected significantly varied genes in CMAP data by setting a threshold of absolute z-scores (Methods), and these genes consist of two equal parts: up-regulated genes (genes with positive z-scores) and down-regulated genes (genes with negative z-scores). Model performance was evaluated using direction accuracy (ACC), calculated by comparing the predicted results to the ground-truth labels from CMAP data (Methods). In total, 21,361 perturbations across the six cell lines were included in this benchmarking analysis (**Fig. 4c**).

We included several fundamental baselines as benchmarking methods, including the Correlation method and DirectTransfer method (Methods). The Correlation method used PCCs among genes as associated change factors for *in silico* perturbation; when a gene is perturbed, genes with higher PCCs relative to it are more significantly affected (Methods). For DirectTransfer, we utilized two sources: genome-scale Perturb-seq data on K562 cells (DirectTransfer_K562), and from the L1000 platform^26^, where PC3 data was selected as the transfer reference as it showed a high number of perturbations (DirectTransfer_L1000). Additionally, we benchmarked deep learning methods such as CellOracle, CPA, scGPT, and scFoundation, which were trained on a merged data collection of scPerturb datasets we curated (Methods). To provide a comparison baseline, a random prediction approach (referred to as Random) was included, where genes are randomly upregulated or downregulated.

We observed a distinctive expression pattern of CMAP’s significant varied genes in single-cell data: up-regulated genes in CMAP tend to exhibit low expression levels in single-cell data, whereas down-regulated genes in CMAP often display high expression levels in single-cell data. As shown in **Fig. 4d** and **Fig. S4**, across all the cell lines, this trend is consistent across all analyzed cell lines. Building on this observation, we introduced an AverageSC method for this scenario, which simply averages the expression values of all landmark genes in single cells to generate predictions. Additionally, we found that single-cell foundation models often predict lower gene expression values relative to the ground-truth values (**Fig. S5**). This bias will complicate the accurate assessment of the model performance. To mitigate this issue, we proposed a strategy for single-cell foundation models that involves using predictions for unperturbed cells as a reference (Methods), aiming to more precisely capture perturbation effects.

We visualized the performance of each method on individual cell lines (**Fig. 4e**) and summarized the overall accuracies for predicting both up-regulated genes and down-regulated genes for all methods (**Fig. 4f**). CellOracle failed to predict the direction of gene expression changes across all the cell lines. Similarly, CPA performed poorly, with results slightly worse than Random in certain cell lines, such as HepG2. The Correlation method showed marginal improvement over Random (**Fig. 4f**). Among the DirectTransfer methods, DirectTransfer_K562 performed at a Random level, while DirectTransfer_L1000 achieved the highest overall accuracy, with a mean value of 0.692. This suggested that transferring gene expression changes from the same platform, such as the L1000 platform, can provide a reliable reference. Both scGPT and scFoundation showed better performance than Random (**Fig. 4f**). Specifically, scGPT performed better than Random in cell line HT29, HepG2, and A549, but performed at Random levels in other cell lines. In contrast, scFoundation outperformed Random in A375, PC3, HT29, and A549. Interestingly, scGPT demonstrated higher accuracy for predicting up-regulated genes, while scFoundation performed better for down-regulated genes (**Fig. 4f**). The AverageSC method achieved the best performance aside from DirectTransfer_L1000, indicating that leveraging the inherent patterns in perturbation data can yield strong predictive results.

Overall, only DirectTransfer_L1000 and the AverageSC method achieved significantly better performance than Random. However, DirectTransfer_L1000 relies on data from the same platform, which is typically unavailable in practical applications, limiting its utility. Meanwhile, the AverageSC method, which merely averages gene expression values, does not account for gene perturbation mechanisms and is therefore impractical for understanding or modeling perturbation effects. Most other methods failed to predict the direction of change for significant genes, highlighting the limitations of current approaches in predicting perturbation effects in zero-shot scenarios. While single-cell foundation models demonstrated some potential to exceed Random performance, their best accuracy on an individual cell line was approximately 0.55—only marginally above random and still far from practical applicability.

We further evaluated the performance of scGPT with varying scales of total perturbations in the merged data collection (Methods), using 10%, 40%, 70%, and 100% of the perturbations. Results indicated a clear scaling effect, wich scGPT achieving progressively higher performance as the number of perturbations increased (**Fig. S6**). We anticipated that as more single-cell perturbation data becomes available across a wider range of cell types and genome-scale gene perturbations, the performance of single-cell foundation models is likely to improve. However, for now, quantitatively predicting perturbation effects in a zero-shot transfer scenario using *in silico* perturbation methods remains a significant challenge.

### Benchmarking in the cell state transition prediction scenario

Many biological processes, such as organ development and cellular reprogramming, are driven by individual genes or small sets of key genes. Evaluating whether computational methods can accurately predict cell state transitions influenced by these genes is critical for assessing their applicability to researchers’ own data and for guiding experimental design.

To establish a benchmarking framework, we collected four single-cell datasets representing distinct biological processes (Methods): Sun2022^29^ for validating that the knockout of *PDCD1* can promote the recovery of exhausted T cells, restoring their effector and memory functions^40, 41^; Ainciburu2023^30^ for validating that SPI1 and *GATA1* perform as a canonical two-gene toggle-switch motif driving the differentiation of hematopoietic cells^6, 42^, where SPI1 is demonstrated to promote the granulocyte-monocyte progenitor (GMP) lineage, and GATA1 commits to the megakaryocyte-erythrocyte progenitor (MEP) lineage; Nair2023^31^ for validating fibroblasts can be reprogrammed into stem cells by four transcription factors (TFs): POU5F1, SOX2, KLF4, and MYC^43^; Steele2020^32^ for validating the knockout of *PTF1A*^44, 45^can induce acinar cells transforming into duct-like cells (acinar-ductal metaplasia, ADM). UMAP visualization of these datasets, along with the ground-truth cell state transitions, was shown in **Fig. 5a**. To evaluate performance, we visualized cell transition trends after *in silico* gene perturbations using streamline plots (**Fig. 5b-f**, Methods).

**Fig. 5.**
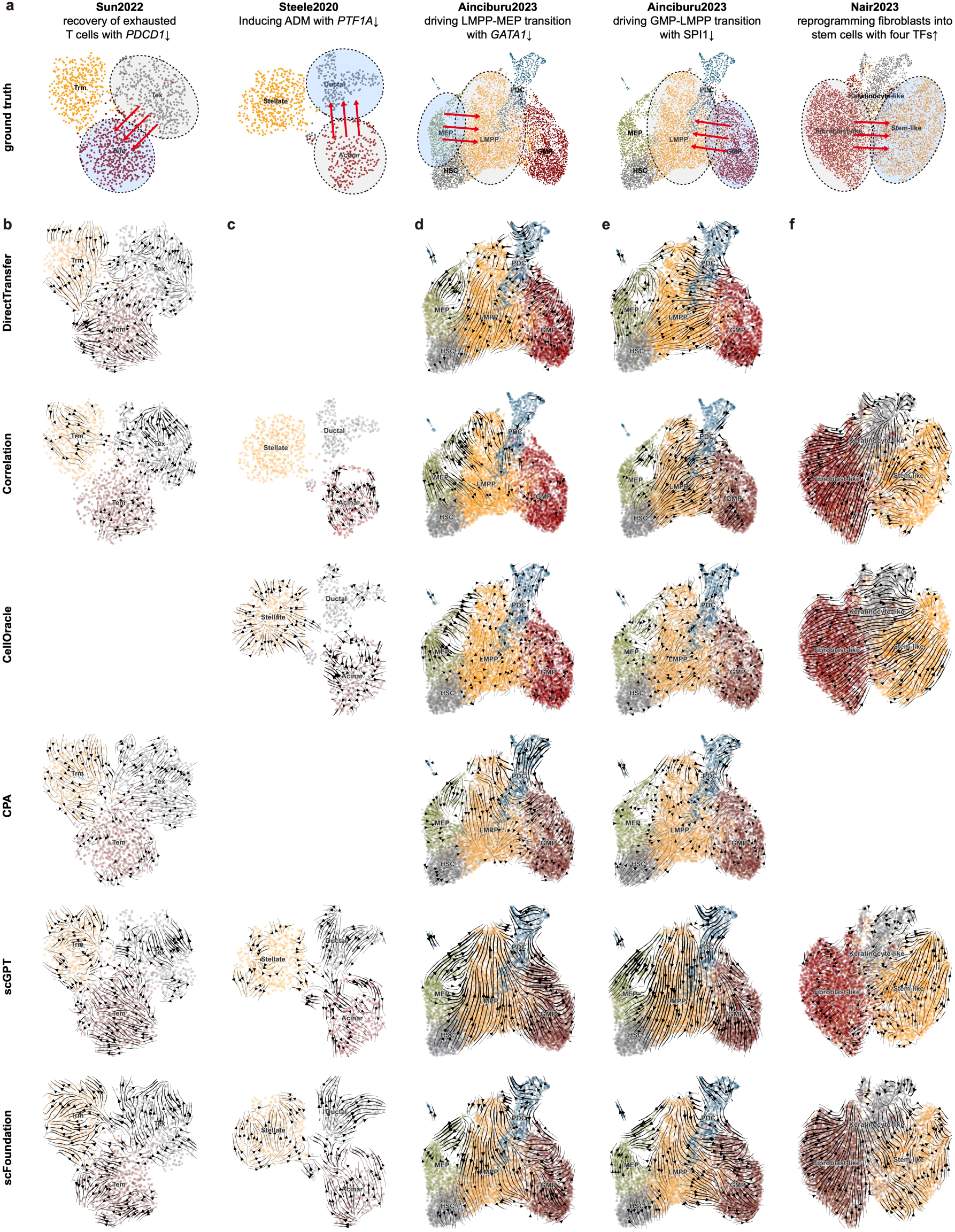
Benchmarking in the cell state transition prediction scenario. **(a),** Ground-truth cell state transitions resulting from the knockdown or overexpression of specific genes. Transition directions are visualized in UMAP space. **(b-f)** Streamline plots display the predicted results from 6 methods across five cases: recovery of exhausted T cells with *PDCD1* knockdown (b), induction of ADM with *PTF1A* knockdown (c), MEP transitioning to LMPP with *GATA1* knockdown (d), GMP transitioning to LMPP with *SPI1* knockdown (e), and fibroblasts reprogrammed into stem cells by four TFs: POU5F1, SOX2, KLF4, and MYC (f).

We benchmarked the methods same as in the zero-shot transfer scenario: DirectTransfer, the Correlation method, CellOracle, CPA, scGPT, and scFoundation. It should be noted that not all methods can be applied to all four datasets. For instance, CellOracle is restricted to *in silico* perturbations of TFs with known binding motifs, and *PDCD1* in the Sun2022 dataset is not included in its TF list. Similarly, CPA and DirectTransfer rely on reference data with the matching perturbations, which were unavailable for the Nair2023 dataset and for PTF1A in the Steele2020 dataset, as these perturbed genes are not included in our filtered scPerturb datasets.

The cell states in the Sun2022 and Steele2020 dataset exhibited discrete cell states, while cells in Ainciburu2023 and Nair2023 form continuous trajectories. Accurately predicting transitions in datasets with discrete cell states requires models to capture more substantial changes in cell states following gene perturbation. As shown in the UMAP plots of both pre-and post-perturbation cells (**Fig. S7**), the predicted gene expression changes are subtle, with cells tending to remain in their original states. In streamline plots, most methods fail to capture the correct transition trends (**Fig. 5b-c**).

In the Ainciburu2023 dataset, both the Correlation method and CellOracle successfully predicted the transitions from MEP to LMPP following *GATA1* knockout, and from GMP to LMPP following *SPI1* knockout (**Fig. 5d-e**). However, other methods failed to capture these specific lineage transitions. In the Nair2023 dataset, CellOracle provided the most accurate visualization of the transition from fibroblast-like cells to stem-like cells in the streamline plot (**Fig. 5f**). scGPT also exhibited an accurate trend in the plot, though it was less pronounced. In contrast, the Correlation method misrepresented the transition, erroneously depicting fibroblast-like cells shifting to keratinocyte-like cells instead of stem-like cells.

Additionally, it is worth noting that CPA failed to predict cell state transitions across all datasets. This limitation may be due to CPA’s reliance on reference data, which can introduce batch effects, resulting in discrepancies between the predicted and actual cell states (**Fig. S7**).

In summary, predicting transitions between discrete cell states remains a significant challenge for current methods. For cell states within continuous trajectories, methods that leverage gene expression correlations, such as the Correlation method and CellOracle, which employed a regression model to infer relationships among genes, perform well in capturing the overall transition trends. Methods that rely on reference training data often struggle to capture accurate cell state transitions, likely due to discrepancies in cell types or experimental platforms between the reference data and the target dataset. Single-cell foundation models, which are trained on large datasets, show promising results in certain cases. For instance, scGPT demonstrates notable performance on the Nair2023 dataset. However, their performance may still be limited by the characteristics of the dataset and the complexity of the transitions being modeled. While single-cell foundation models hold potential, they are not yet universally applicable and require further refinement to achieve broader applicability in diverse scenarios.

## Discussion

Our benchmark categorized *in silico* perturbation methods into four distinct usage scenarios. We observed that no single method consistently excels across all scenarios, highlighting the need of tailoring approaches to specific applications. Single-cell foundation models demonstrated strong performance in predicting unseen perturbations while the DirectTransfer method, despite its simplicity, outperformed others in the unseen cell type transfer scenario. For the cell state transition prediction scenario, methods like the Correlation method and CellOracle proved to be the most effective. In some cases, although baseline methods achieved good performance, they may not be practically useful. For instance, DirectTransfer completely overlooks the variability across different cell types, failing to offer new insights. These findings underscore the importance of aligning method development with the intended usage scenario. Clearly defining the applicable scenarios for each method can not only guide their improvement but also help researchers select the most suitable approach for their specific needs.

One of the most appealing aspects of *in silico* perturbations is that researchers can easily apply them to their data of interest and conduct virtual perturbation experiments. However, due to current data limitations, most usage scenarios of *in silico* gene perturbations still rely on zero-shot predictions, where no training data is provided. Our benchmarking results reveal that existing methods barely outperform Random predictions in terms of quantitatively predicting gene expression changes in zero-shot predictions.

Although achieving reliable predictions with *in silico* perturbation methods remains a long-term challenge, advancements in this field offer hope for future progress. With the ongoing development of biological technologies, single-cell perturbation experiments are expected to cover a broader range of cell types and perturbations. The gradually accumulation of such data will likely enhance the performance of computational methods. Rood el al.^46^ proposed a concept of “perturbation cell atlas”, but how to curate and build such an atlas remains open challenges. It is also important to note that due to the destructive nature of gene perturbation experiments and the technical noise inherent in single-cell sequencing methods, existing data often contains significant levels of noise. As a result, some experimental data may not serve as reliable gold standards. Given these uncertainties, integrating computational predictions with biological experiments in a closed-loop approach could potentially accelerate our understanding of gene expression changes and their impact on cellular phenotypes. Beyond data, innovation in model architectures is also essential. Current transformer-based models show promise in leveraging massive datasets, but there is considerable potential for exploring alternative architectures. For instance, recent studies have highlighted the potential of diffusion models for performing *in silico* perturbations^47^, offering a promising direction for further research and development in this domain.

While experimental data can cover individual perturbations at the genome scale on a single cell type, extending these experiments to cover all possible perturbation combinations remains infeasible. *In silico* methods present a promising solution to guide the selection of perturbation combinations for experimental validation. Several studies have attempted to benchmark *in silico* methods for predicting the effects of perturbation combinations^8, 9, 18, 48^. However, we believe that the currently available single-cell perturbation data involving simultaneous perturbation of multiple genes is insufficient for a comprehensive benchmarking. With the accumulation of more extensive and diverse data in the future, we plan to incorporate this important benchmark to address the growing need for evaluating combination perturbation predictions.

Our benchmark focuses exclusively on *in silico* methods designed for scRNA-seq data, as it is the most abundant resource for single-cell perturbation studies and serves as the foundation for the development of most existing methods. Single-cell perturbation methods for sequencing other modalities, such as chromatin accessibility^49, 50^ and protein abundance^51^, have also been developed. We anticipate that as datasets for these modalities continue to grow, *in silico* methods capable of predicting molecular profiles beyond RNA expression will emerge. These advancements could enable predictions of chromatin states, protein abundance, and other intracellular molecular features, paving the way for a deeper understanding of cellular systems and facilitating the design of more precise and robust perturbation experiments.

To support researchers in evaluating methods with our benchmarking framework, we have provided a comprehensive pipeline in our GitHub repository: https://github.com/Chen-Li-17/CellPB. We developed a Python module CellPB for users to easily use our benchmark. For each usage scenario, we implemented intuitive and flexible interfaces that allow seamless access to benchmarking datasets and straightforward computation of benchmarking metrics. Additionally, we included examples of each *in silico* perturbation method benchmarking in this study, offering clear guidance for training and evaluating their own models. These resources aim to streamline the benchmarking process and promote method development. We believe this work will serve as a foundation for advancing *in silico* perturbation methods and will contribute to progress in the field of biomedical research.

## Methods

### Data curation

#### scPerturb data

We utilized the scPerturb scRNA-seq datasets (version 1.3, available at https://zenodo.org/records/10044268) as benchmarking data for our study. scPerturb is a comprehensive resource of single-cell perturbation datasets, encompassing a wide range of perturbations, including gene modifications, small molecules, cytokines, and more. This dataset includes a total of 50 RNA and protein datasets. To ensure consistency and relevance, we excluded datasets that do not map CRISPR perturbations to specific genes and those derived from non-human cells.

After applying these filtering steps, we retained 17 single-cell perturbation datasets for analysis, each representing cells of a specific cell type perturbed by various genes. To prepare the data for model training and evaluation, we performed the following preprocessing steps:

- Feature Selection: We identified the top 5,000 highly variable genes (HVGs) from each dataset using standard procedures. We also included all perturbed genes originally present in the dataset’s feature list.
- Normalization: Raw count data was normalized using the log-normalization method, implemented through the standard preprocessing workflow of Scanpy^52^.
- Control-cell pairing: For every cell subjected to perturbation, we randomly sampled a control cell from the same dataset to form a pre-and post-perturbation cell pair. These pairs served as the basis for training and evaluating models.
- Perturbation filtering based on cell thresholds: Perturbations with fewer than 100 cells were excluded to ensure robust analysis with sufficient data.
- E-distance calculation: The E-distance for each perturbation was computed to quantify the perturbation’s effect. E-distance is a statistical measure of distance between two distributions, which is utilized to indicate the degree of perturbation effect in scPerturb. We utilized the scPerturb Python library to compute this metric between control cells and perturbed cells. For each perturbation, we sampled 200 cells and selected 2,000 HVGs, as instructed by the scPerturb documentation, to perform the calculation.
- sgRNA effectiveness filtering: To address variability in sgRNA efficacy, we retained only sgRNAs that demonstrated consistent perturbation effects. Specifically, if an sgRNA’s E-distance exceeded 10 and was at least 10-fold greater than another sgRNA targeting the same gene, this sgRNA was included, while the underperforming sgRNA was excluded.
- Identifying differentially expressed genes (DEGs): For each perturbation, we identified DEGs by comparing perturbed cells to control cells. Genes were ranked based on adjusted *p*-values computed using the Wilcoxon test in Scanpy, with those having adjusted *p*-values < 0.05 designated as DEGs.
- Data splitting for training, testing, and validation: The filtered perturbations were divided into training, testing, and validation sets in a 7:2:1 ratio. The training set was used to fit the models, the validation set to select the best model, and the testing set to evaluate the final model performance.

#### CMAP data

CMAP data is based on L1000, a high-throughput gene expression profiling technology that converts raw fluorescence intensity of genes into expression levels. CMAP data contain over 1.5 million gene expression profiles derived from approximately 3,000 gene perturbations and 5,000 drug perturbations across multiple cell lines. For each perturbation in each cell line, CMAP provides a normalized z-score vector of 978 landmark genes, representing the extent to which these genes are affected by the perturbation in the given cell line. This dataset is a valuable resource for understanding gene expression changes induced by perturbations.

The CMAP data is organized into five distinct levels, each representing a different stage of data processing: Level 1 records the raw fluorescence intensity measurements. Level 2 and level 3 provide deconvoluted and inferred gene expression values, respectively. Level 4 provides robust z-scores derived from the expression data. Level 5 aggregates the z-scores, offering a comprehensive summary of perturbation effects. In our study, we used level 5 data as the benchmark ground truth due to its aggregated and robust representation of gene expression changes.

To align the CMAP data with single-cell datasets, we collected single-cell data with six overlapping cell lines: A375, PC3, MCF7, HT29, HepG2, and A549. These single-cell datasets were used as control cell inputs for model evaluation. Each single-cell dataset underwent the following preprocessing steps: (1) 3,000 HVGs were selected, with perturbation genes from the CMAP data added to the feature set, and (2) raw count data was log-normalized.

For each cell line, we calculated the PCC between the single-cell data and the CMAP data. For the single-cell data, we selected 3,000 HVGs from the single-cell data and averaged the expression levels across all cells to obtain a mean expression vector. For the CMAP data, we extracted control samples corresponding to the specific cell line and averaged them to a mean expression vector with a length of 978. The PCC was then computed using the genes common to both mean expression vectors.

For each perturbation in the CMAP dataset, the level 5 data provide an aggregated z-score vector representing the gene expression changes. Positive z-scores indicate increased expression following the perturbation (up-regulated), while negative z-scores indicate decreased expression (down-regulated). To refine the analysis, we selected genes with z-scores greater than 1.5 or less than –1.5 as significantly affected and excluded genes with an expression level less than 0.1 in the single-cell datasets. From this filtered gene set, we further sampled genes to ensure an equal representation of positive and negative z-scores, balancing the directionality of expression changes. These genes were identified as top significant genes, and we calculated direction accuracy using these genes to evaluate model performance.

#### Single-cell data with cell state transitions

We compiled four single-cell datasets encompassing five distinct biological processes for our study. To ensure consistency and minimize batch effects, we selected data from a single donor for datasets with multiple donors.

- Sun2022 dataset: This dataset originates from cells within the tumor microenvironment of gastric cancer. We selected donor GC05 and retained three CD8+ T cell subsets: exhausted CD8+ T cells (Tex), effector memory CD8+ T cells (Tem), and residual memory CD8+ T cells (Trm). The biological process of interest involves the knockdown of *PDCD1*, which is expected to induce a transition from Tex to Tem.
- Ainciburu2023 dataset: Derived from hematopoietic cells in a human cell atlas, this dataset includes data from the 19-year-old donor with the following cell types: lympho-myeloid primed progenitors (LMPPs), granulocyte-monocyte progenitors (GMPs), megakaryocyte-erythrocyte progenitors (MEPs), hematopoietic stem cells (HSCs), and plasmacytoid dendritic cells (pDCs). Two biological transitions were investigated: knockdown of GATA1 leading to MEP-to-LMPP transition, and knockdown of SPI1 leading to GMP-to-LMPP transition.
- Nair2023 dataset: This dataset originates from a fibroblast reprogramming experiment. We included three cell types for analysis: fibroblast-like cells, keratinocyte-like cells, and stem-like cells. The focus is on the reprogramming of fibroblast-like cells into stem-like cells via the overexpression of canonical reprogramming factors: POU5F1, SOX2, KLF4, and MYC.
- Steele2020 dataset: This dataset was obtained from a human pancreas cell atlas, using data from donor N1. We selected acinar cells, ductal cells, and stellate cells as cell types of interest. The biological process studied is Acinar-Ductal Metaplasia (ADM), a transition from acinar cells to ductal cells, which can be induced by the knockdown of *PTF1A*.

All datasets were processed using a standardized pipeline, which included log-normalization of the count data followed by the selection of HVGs using Scanpy with parameters set to min_mean=0.0125, max_mean=3, and min_disp=0.4.

### Implement of *in silico* perturbation methods

In this study, we evaluated 10 *in silico* perturbation methods across four distinct benchmarking scenarios. Some of the methods and applicable to multiple scenarios. For each method, parameter settings were fixed according to official guidelines and tailored to align with our benchmarking datasets.

#### Linear

The linear method is adopted from another benchmarking study^22^. The optimization objective is

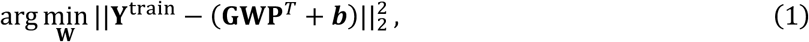

where **Y**^train^ is the gene expression matrix of perturbed cells, rows represent the gene feature set, and columns represent the perturbed genes in the training data. **G** is the gene embedding matrix, which is obtained from the top *k* principal components from the PCA (principal component analysis) on **Y**^train^. **P** is subset from **G** with the perturbed genes in the training data. **W** ∈ ℝ*^k^*^×*k*^ is the fitted matrix, and ***b*** is the vector of average gene expressions of 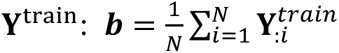
. **W** is solved using the normal equations:

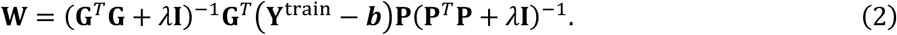

Here, we set *k* to 512 and *λ* to 0.1. Then the fitted W is used for prediction:

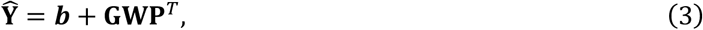

and **Y̑** is the predicted gene expression, and **P** subset from **G** denotes the embeddings of perturbed genes in the testing data.

#### Linear with scGPT embeddings

Linear-scGPT employs the same modeling approach as the linear method, with the key distinction being that the gene embedding matrix **G** is derived from the scGPT model rather than the PCA components of the training data. We followed the instruction from GitHub repository of the Linear method to obtain the scGPT gene embeddings: https://github.com/const-ae/linear_perturbation_prediction-Paper/blob/main/benchmark/src/extract_gene_embedding_scgpt.py.

#### Correlation

We first constructed a gene-gene correlation matrix, where each element represents the PCC between two genes. When a gene is perturbed, the expressions of other genes change according to a scaling factor determined by the corresponding PCCs. If a gene is knocked down, its expression is set directly to 0. If a gene is overexpressed, 1 is added to its original normalized expression value.

#### CellOracle

We followed the tutorial provided on the CellOracle website at https://morris-lab.github.io/CellOracle.documentation/tutorials/simulation.html. For each dataset, the GRN was inferred from control cells and subsequently applied to simulate expected gene expression changes under gene perturbations. Since scATAC-seq data from the same tissue as the scRNA-seq datasets is unavailable, we utilize the function ‘load_human_promoter_base_GRN’ to provide base GRNs for all datasets.

#### scGen

We followed the instructions provided on the scGen website via https://scgen.readthedocs.io/en/stable/tutorials/scgen_perturbation_prediction.html. For each perturbation transferred to an unseen cell type, we train a model for prediction, similar to the procedures used for CPA and CellOT.

#### CPA

We followed the tutorial provided on the CPA website: https://cpa-tools.readthedocs.io/en/latest/tutorials/Kang.html. While CPA was originally designed for the unseen cell type transfer scenario, we adapted it for application in zero-shot transfer and cell state transition prediction scenarios. In these cases, the training data was sourced from the scPerturb datasets.

#### CellOT

We adopted the codes provided on the CellOT GitHub repository: https://github.com/bunnech/cellot. Since our control group data contains significantly fewer cells compared to the tutorial dataset, we reduced ‘n_iters’ from 100,000 to 1,000 to improve computational efficiency.

#### GEARS

We followed the tutorials provided on the GEARS Github repository: https://github.com/snap-stanford/GEARS. GEARS is designed for the unseen perturbation scenario. We first reformatted our data to align with GEARS’ input requirements. The model was then trained using its default parameter settings, with the number of epochs to 20. The best-performing model, selected based on its performance on the validation dataset, was used to evaluate the test dataset.

#### scGPT

The scGPT model is versatile in various scenarios for its ability to accept any length of genes as input. In unseen perturbation transfer and unseen cell type transfer scenarios, we followed the guidance on scGPT Github repository to reformat data and train models: https://github.com/bowang-lab/scGPT/blob/main/tutorials/Tutorial_Perturbation.ipynb. Default parameter settings were used, except that the model was trained with all genes in the feature set. Similar to GEARS, we trained the scGPT model for 20 epochs and selected the best-performing model based on validation results for testing. For zero-shot transfer and cell state transition scenarios, the scGPT model was trained on a combined dataset comprising a total of 17 datasets, ensuring broader generalization across different biological contexts.

#### scFoundation

scFoundation also employs transformer-based architectures, making it suitable for *in silico* perturbation tasks, similar to the training paradigm of scGPT. We adopted the code from scFoundation Github repository: https://github.com/biomap-research/scFoundation. Specifically, we leveraged the encoder module of the scFoundation model to train the perturbation model. The training and testing procedures followed the same setup as in scGPT. Given that scFoundation incorporates more transformer layers than scGPT, we set the learning rate to 1e-5.

#### Training and application of single-cell foundation models for the zero-shot transfer scenario

We curated 17 processed scPerturb datasets to serve as the training data for scGPT and scFoundation. Each dataset was carefully processed and split into training, testing, and validation subsets. These subsets were subsequently merged across all datasets to create comprehensive training, testing, and validation datasets. The models were trained on the combined training set using random shuffling to ensure robust learning. Considering that the cells in the same batch may come from different datasets and hold gene features with different lengths, we performed the data padding with ‘MASK’ token, which are not involved in training models. For both scGPT and scFoundation, we trained the model for 20 epochs and selected the best-performing models as the final model for zero-shot transfer.

We observed that when trained single-cell foundation models are applied to novel datasets, the predicted gene expression values tend to be lower than the ground-truth gene expression values (Fig. S5). To address this bias, we employed a correction strategy using control cells. Specifically, let the gene expression of control cell *i* be denoted as *x_i_*., and the model’s predicted output with perturbation as *y_i_*. We then used the model to predict the gene expression of the same control cell without perturbation, denoted as *y_i_*^’^. The difference *y_i_*_*i*_− *y_i_*^’^ was taken as the final predicted gene expression change.

### Benchmarking metrics

We utilized different metrics in different scenarios to comprehensively and fairly evaluate all the *in silico* perturbation methods. In unseen perturbation transfer and unseen cell type transfer scenarios, the single-cell perturbation datasets from scPerturb are adopted for model training and testing. For each perturbation, we denote *x*_*i*_ ∈ ℝ*^n^* and *y*_*i*_ ∈ ℝ*^n^* as average gene expression vectors of control cells and perturbed cells, respectively, and *n* denotes the length of gene feature set.

#### Pearson correlation coefficient (PCC)

The PCC value is calculated by the following equation:

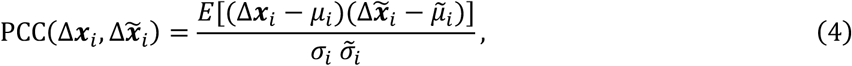

where Δ***x_i_*** and Δ***x̃_i_***. represent the ground-truth and predicted changed expression vectors of perturbation *i*, respectively. Here, *μ*_*i*_ and *μ̃_i_* denote the mean values of Δ***x***_*i*_ and Δ***x̃_i_***, while *σ*_*i*_ and *σ̃_i_* are their corresponding standard deviations. PCC values are computed specifically for DE genes in each perturbation.

#### Direction accuracy (ACC)

We calculated ACC by the following equation:

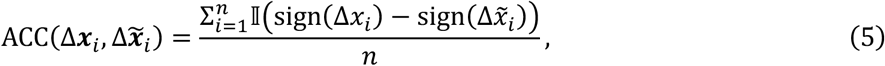

where ‖(⋅) is an indicator function defined as:

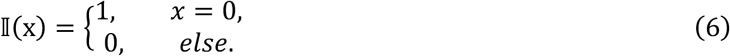

In the unseen perturbation transfer and unseen cell type transfer scenarios, ACC values are calculated for DEGs. While in the zero-shot transfer scenario, the ACC values are computed for the top significant genes from the CMAP dataset.

#### Mean absolute error (MAE)

The MAE is calculated for the non-DEGs, including stably highly expressed genes (SHGs) and stably lowly expressed genes (SLGs) for each perturbation, and it is defined as the following equation:

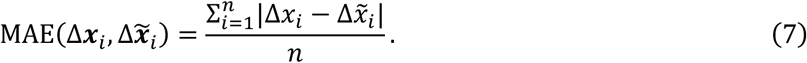

#### Streamline plot

The model’s predicted expression change vector was used as the cellular velocity for each cell. Subsequently, the ‘velocity_embedding_stream’ function from the scVelo^53^ package was employed to visualize the streamlines on the UMAP plot.

## Data availability

The scPerturb datasets were downloaded from https://zenodo.org/records/10044268. The CMAP data was downloaded from Gene Expression Omnibus (GEO) with accession number <u>GSE92742</u>. The scRNA-seq datasets containing 6 cell lines intersected with CMAP data were obtained from the Broad Institute’s single-cell portal (<u>SCP542</u>) and Sequence Archive (CNSA) with accession number <u>CNP0003658</u>. scRNA-seq datasets used in the cell state transition prediction scenario were downloaded from different resources: Sun2022 dataset (<u>OMIX001073</u>), Ainciburu2023 dataset (<u>GSE180298</u>), Nair2023 dataset (<u>GSE242424</u>), and Steele2020 dataset (<u>GSE155698</u>).

## Code availability

Our code for benchmarking is available at https://github.com/Chen-Li-17/CellPB.

## Acknowledgements

The work is supported in part of National Natural Science Foundation of China (62250005, 62373210, 62433001, and 92470105) and The National Key R&D Program of China (2021YFF1200900). Figure 1 was created with BioRender.com.

## Author contributions

C.L., H.G. and X.Z. conceived and designed the study. X.Z, K.L and Y.S. supervised the study and provided critical revisions. C.L. and H.G. collected the data. C.L. implemented the methods, conducted the experiments, and analyzed the results. C.L. and L.W. drafted the manuscript and prepared the figures. X.Z., K.L., H.G., and Y.S. reviewed the manuscript and provided critical insights. All authors contributed to reviewing and approving the final manuscript.

