## Supplementary figures for "Benchmarking AI Models for *In Silico* Gene Perturbation of Cells"

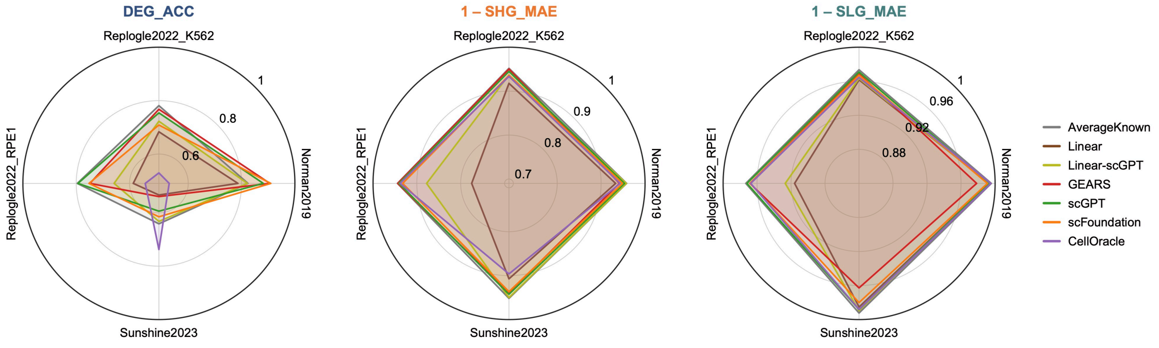


**Supplementary Fig. 1.** Radar plots of DEG_ACC, SHG_MAE, and SLG_MAE for 7 methods across 4 selected datasets.

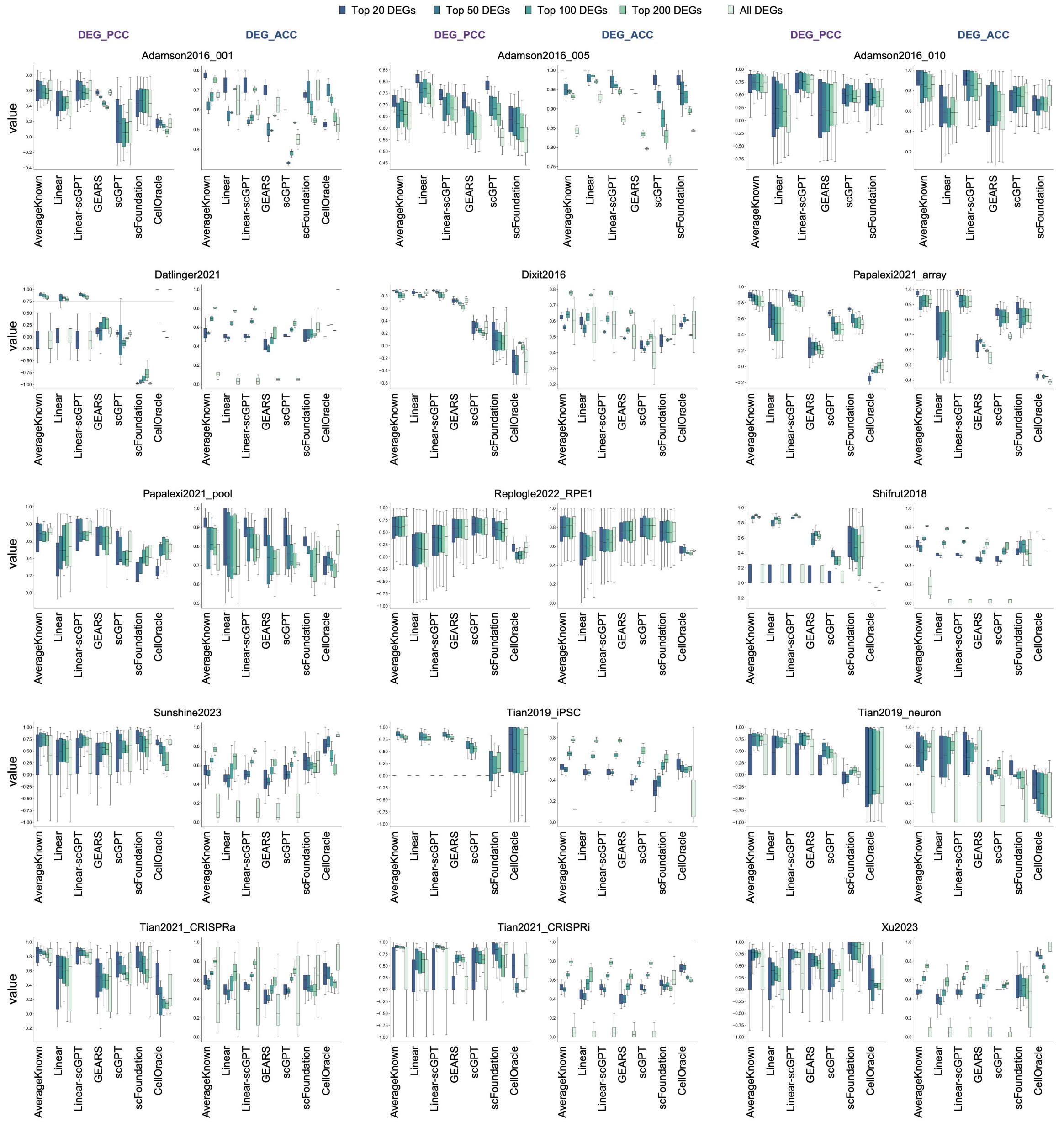


**Supplementary Fig. 2.** Box plots comparing DEG_PCC and DEG_ACC of 7 methods across 15 single-cell perturbation datasets.

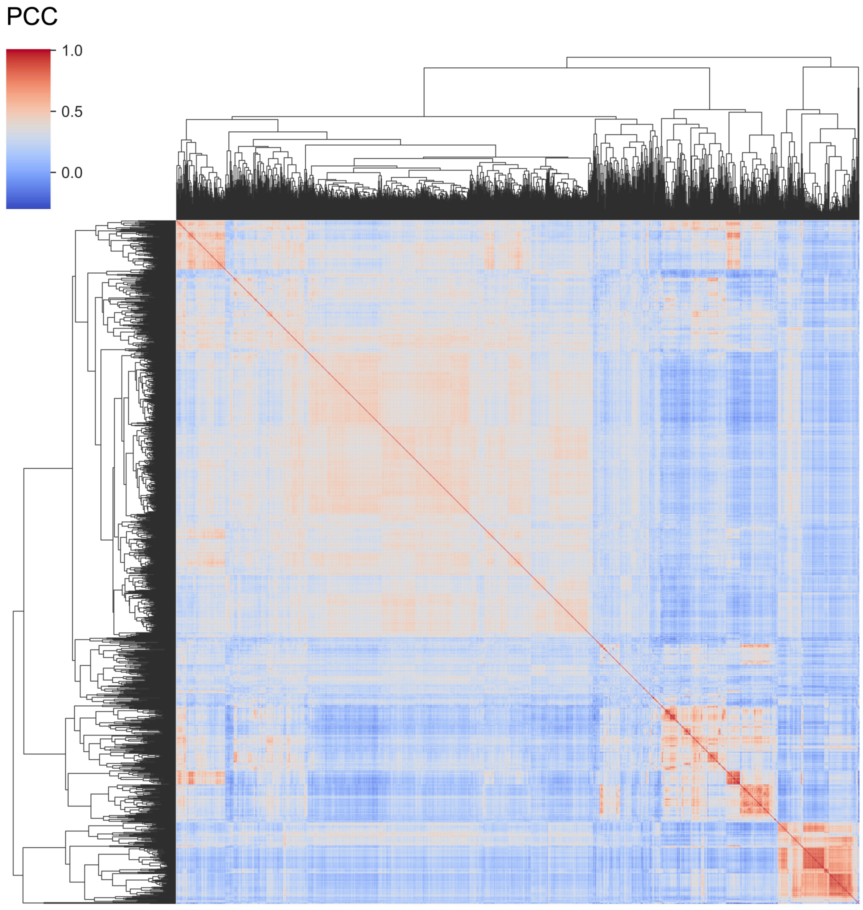


**Supplementary Fig. 3.** Hierarchically-clustered heatmap of correlation matrix among Replogle2022_K562's 1,299 perturbations.

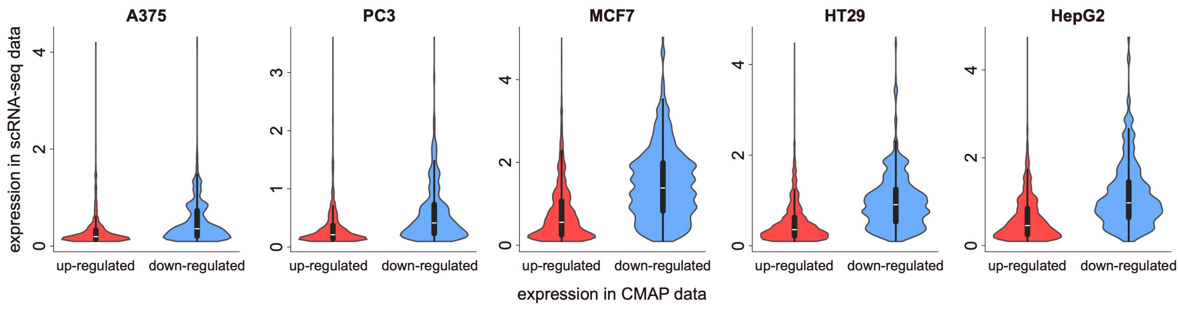


**Supplementary Fig. 4.** Expression levels of CMAP's up-regulated and down-regulated genes in single-cell data.

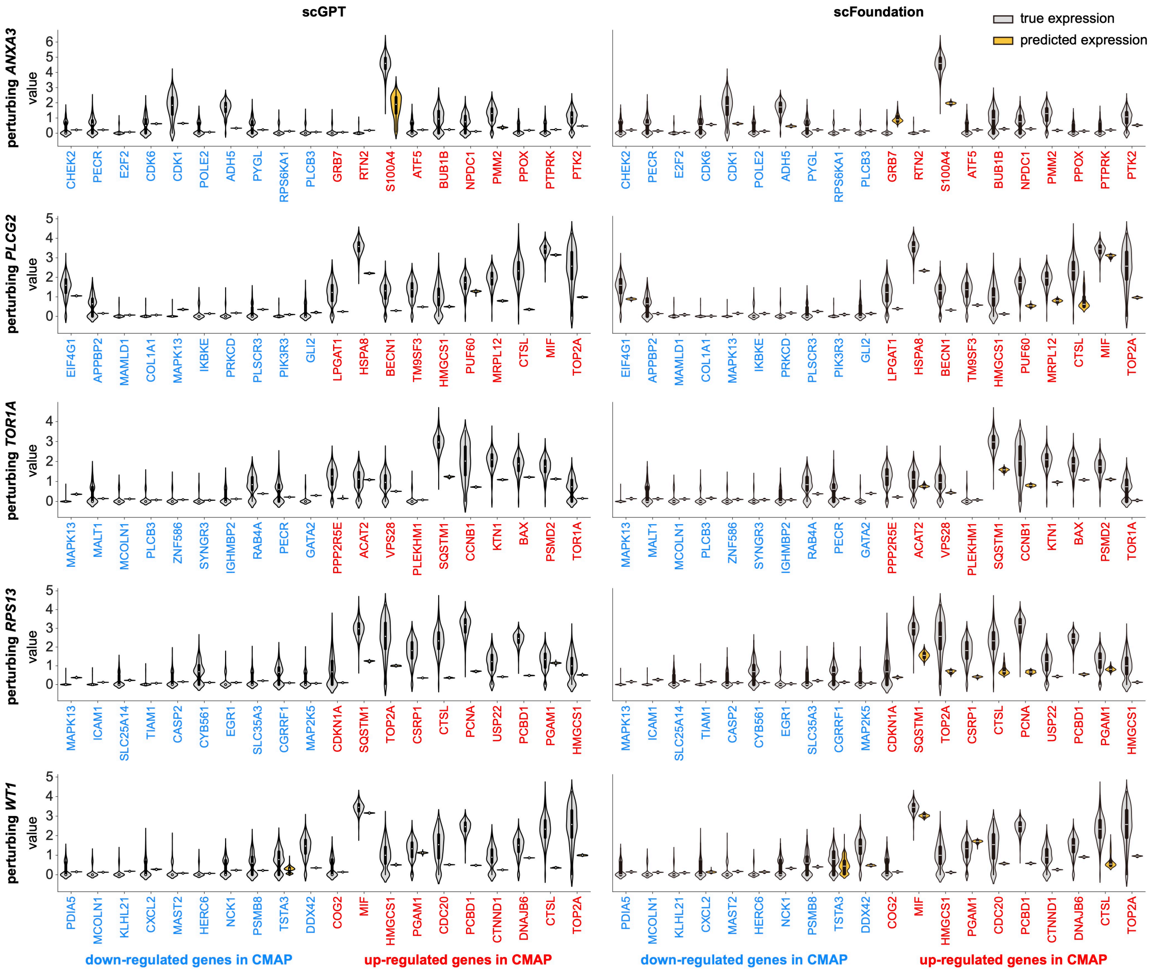


**Supplementary Fig. 5.** Comparison of ground-truth and predicted gene expression of single-cell foundation models. We randomly selected 5 perturbations (*TOR1A*, *WT1*, *ANXA3*, *RPS13*, and *PLCG2*) in cell line A549 to conduct the comparison. For each single plot, the up-regulated and down-regulated genes are annotated in blue and red, respectively. The left column and right column represent results of scGPT and scFoundation, respectively.

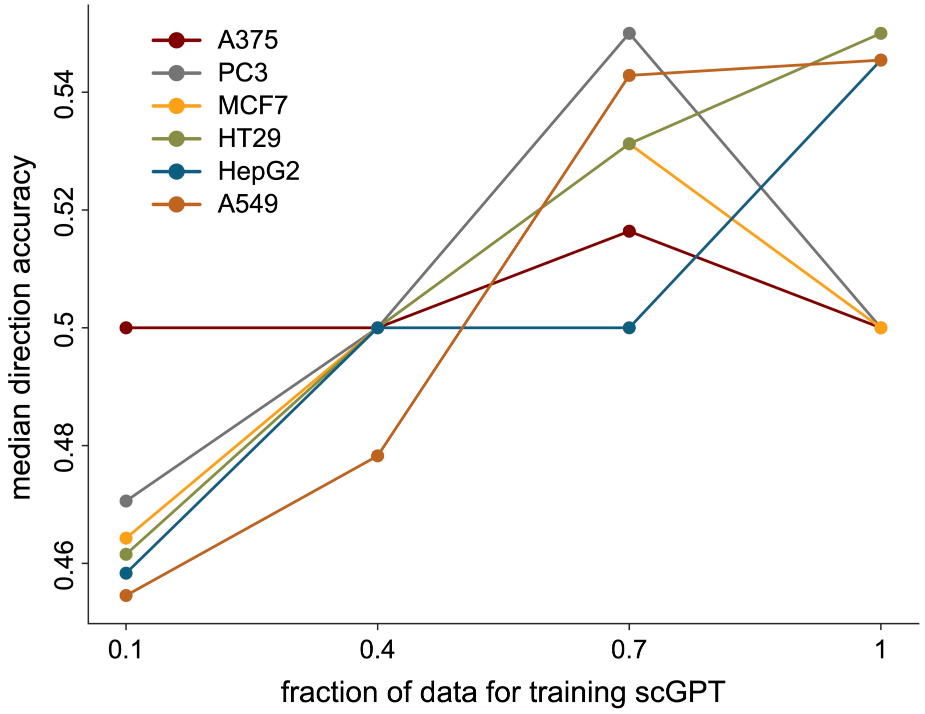


**Supplementary Fig. 6.** scGPT performance at varying data scales in zero-shot prediction scenario. The merged scPerturb data is subset to 10%, 40%, 70%, and 100% for training scGPT, respectively.

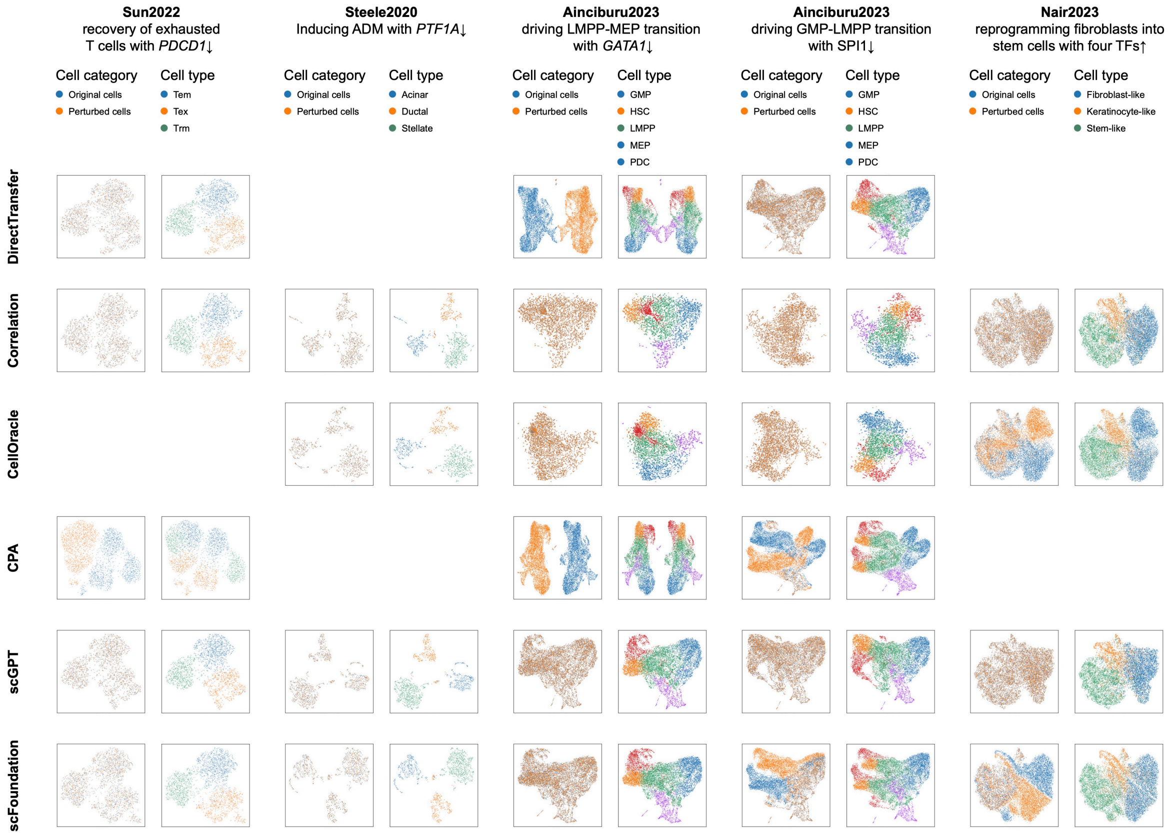


**Supplementary Fig. 7.** UMAP visualization of predicted cells along with original cells. The results of 6 methods across 4 datasets are shown. For each method in each dataset, the cells are annotated with cell type and category (ground-truth or predicted cells).
